# *In silico* Study of the Uncertainly Significant *VCP* Variants reveal major Structural, Functional fluctuations leading to potential Disease-Based Associations

**DOI:** 10.1101/2023.11.04.565542

**Authors:** Tathagata Das, Saileyee Roychowdhury, Parimal Das

## Abstract

Valosin containing protein is involved in a plethora of crucial functions from proteostasis, stress granule clearance to genome maintenance and ubiquitination. Hence, mutations in *VCP* can lead to a plethora of fatal diseases like Amyotrophic Lateral Sclerosis, Inclusion body myopathy with Paget disease of bone and frontotemporal dementia type 1, Spastic paraplegia, Charcot-Marie-Tooth disease type 2Y, Dementia, and Osteitis Deformans to name a few. Studies on *VCP*’s disease phenotype relationship, structural, and functional modifications of the protein’s stability, conservation, molecular dynamics, and post-translational modifications haven’t been performed. This *in silico* study investigated the variants of *VCP* (R95C, R95G, A160P, R191P, R191Q) which have conflicting interpretations of pathogenicity which are often depreciated and lack data. Additionally, this study screens the study cohort and the fatal diseases linked to all these variants. Interestingly through various computational tools and disease-based population studies, it was found that these variants are often found in patients linked with fatal diseases. The protein-protein interaction showed *UFD1* has a direct association with *VCP*. The physicochemical parameters showed that A160P had the highest fluctuations of all the variants. *VCP*’s protein secondary structure, molecular dynamics simulations of RMSD, RMSF, RoG, hydrogen bonds, and solvent accessibility were all comparatively impacted due to the changes caused by the variants. Box-plot and Principal component analysis of the MD simulations visualized the changes in the wild type of the protein. Research in wet labs and screening of patient cohorts is necessary to further characterize the diseases linked with these variants. This can potentially lead to the identification of biomarkers for fatal rare diseases.

## Introduction

The Valosin-containing protein is a hexameric protein of the ATPases associated with the diverse cellular activities (AAA) family, it has critical functions in the quality control of the protein and signalling pathways. It has roles in a plethora of cellular physiology and is directly connected to various other proteins and their functioning. The *VCP*/p97 system has been deemed as an important factor in the ubiquitin system and intracellular pathways. It has its role in mitosis, involved in the fragmentation and reassembly of the Golgi stacks, which is the link to its involvement in the making of the endoplasmic reticulum[1]. The concentration of the protein was found to be 34 micrograms/litre via mass spectrometry[2]. The protein is additionally involved in stalled replication forks, and DNA damage response, where it is called into action for double-stranded breaks and limits the mutations occurring due to DNA damage, along with that it is the negative regulator of the production of type I interferon. Due to the involvement of *VCP* in all the above-mentioned roles it has a direct effect on genome stability and degenerative diseases[3].

Missense mutations in the *VCP* can cause various diseases, like Inclusion body myopathy with Paget disease of bone and frontotemporal dementia type 1, Amyotrophic lateral sclerosis, Spastic paraplegia, Frontotemporal dementia and/or amyotrophic lateral sclerosis 6, Charcot Marie tooth disease, Inclusion body myopathy with Paget disease of bone and frontotemporal dementia, etc[1,4,5]. These recorded diseases fall in the range of “pathogenic” or “likely pathogenic” missense variants in dbSNP. However, disease information is either unavailable or obscure in Clinvar for variants having clinical significance as “Pathogenic likely Pathogenic” or “Conflicting interpretations of pathogenicity”[6]. The frequency of the variants in the reported studies was only screened in small cohorts with rs1554668805 having no frequency data in dbSNP[7]. With the growing number of missense variants in proteins, it should be known which mutations are causing the particular phenotype leading to degenerative diseases. The most prevalent method to confirm and cross-confirm variations is sequencing. With advancements in sequencing techniques, number of variants is growing rapidly. Characterization of novel variants into pathogenic or benign is also necessary. But there is a plethora of variants which have no transparent classification of being fatal or benign. Hence this study focuses on giving a prominent profile to *VCP* variants that fall under uncertain significance of pathogenicity such that during wet lab research these variants can be classified and, further be used as biomarkers for certain fatal diseases[8].

*In silico* tools provide data on conserved regions throughout evolution, structure, protein-protein interactions, favourable conformations, ligand binding sites, location, and function of the proteins to name a few. Gain and loss of function mutations are also predicted through the stabilization energy of the protein and are highly relatable if the conformational energy is changing due to a mutation. There were about 7979 variations reported of *VCP* at the time of this study of which 414 variations were missense variants and 18 were missense-pathogenic. It was identified that out of the total 7979 variants, 2 fell under missense-pathogenic likely pathogenic with inconsistent pathogenicity profile, and 1 variant was under conflicting interpretations of pathogenicity, which had uncertain clinical significance. Several bioinformatic tools were utilized to study the interactions, pathogenicity, and stability of the protein. The physicochemical, structural, functional, and evolutionary features were studied and analysed of the variants classified as pathogenic likely pathogenic and conflicting interpretations of pathogenicity[6]. Diseases associated with *VCP* were scrutinized through various disease and variant databases. It was observed that rs121909332 had a total of 9 SNPs and 17 variant disease association evidence, for rs121909334, 2173 multiple diseases associated SNPs were found. 46 variant disease association with strong evidence of R191Q causing autosomal dominantly inherited ALS, Inclusion Body Myopathy with Paget’s disease of bone and Frontotemporal Dementia (IBMPFD)[9], Paget disease of bone, familial ALS, FTLD-TDP type IV, and Braak stage five Parkinson’s disease was reported [1,4,10,11], rs1554668805 (A160P) interestingly had no record and publications with evidence in NCBI or other resources[6].

## Materials and Methods

### Dataset Preparation

The NCBI dbSNP database was screened to obtain the variants of *VCP*. A total of 7979 results were obtained without any filter. Upon putting filter for only missense mutations 414 rsID results were obtained, benign-missense mutations had 4 results, likely benign-missense gave 8 results, pathogenic-missense gave 18 results, likely pathogenic-missense was 12, pathogenic likely pathogenic-missense had 2 rsID in it and only 1 result was obtained for conflicting interpretations of pathogenicity which fell under missense classification[7]. In this *in silico* research rsID of pathogenic-likely pathogenic and conflicting interpretations of pathogenicity variants were considered. The deleterious nature of the *VCP* variants were investigated through several bioinformatics tools and algorithms. rs121909332, rs121909334, and rs1554668805 were analyzed here and the transcript isoforms were filtered out. Hence, in total 5 amino acid changes were scrutinized[6].

### Protein-Protein Interaction and Disease Pathology of *VCP*

The PPI study presented the empirical data on the interaction networks of *VCP*. Hence to study this protein atlas database was used. Additionally, the pathology of the protein was studied to detect its expression in various diseases. The human protein atlas was used to focus on the expression profiles in human tissues and proteins[12]. The pathology information was based on mRNA and protein expression data from around 17 different forms of human cancers to find the precise marker where *VCP* is involved[13].

### Pathogenicity Analysis

Algorithm-based tools were used to determine, and predict the nature of pathogenicity of the *VCP* variants. The tools used were SIFT[14], Polyphen2[15], Panther[16], SNP&GO[17], PhDSNP[18], MetaRNN[19], Mutpred2[20]. These tools require the wild-type protein sequence and the specific amino acid change site, then it gives the pathogenicity level or score of the specific variant. More details can be studied from [21].

### Protein Stability Analysis

A plethora of tools were used to understand the stability profile of the variants. Gibbs free energy-based algorithms were used like Maestro[22], Cupsat (Cologne University protein stability analysis tool)[23], mCSM (mutation cutoff scanning matrix)[24], I-mutant[25], SDM[26], Dynamut[27], MUPro[28], DUET[24]. All these tools provided ddG values with the score determining whether the *VCP* variants in consideration are stabilizing or destabilizing. Further methodology details can be found in [29].

### Evolutionary Conservation

The Consurf web server was used to study evolutionary conservation. The consurf server is a bioinformatics tool for scrutinizing the evolutionary conservation of amino acids based on phylogenetic relations between homologous sequences. In this study, Uniref90 and Uniprot databases were used separately. The evolutionary conservation of each of the five variants was analysed against the 9-pointer conservation scale (Figure-4) [30]. The evolutionary rate of a particular amino acid is directly dependent on its structural and functional importance. The specific conservation score by position was computed using the Bayesian and Machine Learning algorithms.

### Stereochemical and Physicochemical Analysis

Determination of the secondary structure of the *VCP* protein was done using Psipred workbench [31]. The particular *VCP* variants were analysed using the secondary structure prediction obtained[32]. Discovery Studio was used to induce the mutational changes in the wild-type protein and to draw the Ramachandran plots[33] for each of the five mutations plus the wild-type. All the plots were individually inspected to find whether the variants of concern fell in the allowed or the disallowed regions.

The understanding and determination of the physicochemical properties of the protein aids to decipher the changes of the mutant variants when compared to the wild-type protein. It is necessary to know the physicochemical conditions of the receptor during the design of the ligand drug. Hence to study the physicochemical property of the wild-type *VCP* protein and compare it with the mutant variations ProtScale was used. It detects, computes, and shows the specific profile of an amino acid on the given protein. For this research, the hydropathicity, hydrophobicity, polarity, alpha helix, and beta-sheet plots were drawn for each of the variants, and their respective changes were shown[32]. Expasy Protparam was used for the computation of several physical and chemical parameters like GRAVY value, Instability index, and aliphatic index of the wild-type protein as a reference against the mutants.

### Genome and OMIM Investigation

The UCSC genome browser was used to visualize the genomic data of *VCP*[34]. The involvement of the variants with various diseases was procured and literature survey was done [35] [36]. The Online Mendelian Inheritance in Man was used to gather comprehensive knowledge about the *VCP* and its genetic phenotypes associated with diseases[37]. The three rsIDs of the five variants were used to study the severity of the diseases in association [38].

### Post-Translational Modifications

Post-translational modifications of proteins are important to enhance the functional diversity of the proteins by the addition of various functional groups. It is a necessity to understand the post-translational modifications when analysing the various functionalities of proteins. This eventually puts a transparent view of the causable potential diseases, its prevention and, treatment. For this research, various PTM sites were investigated. Netphos 3.1 (Phosphorylation)[39], NetNGlyc 1.0, NetOGlyc 4.0 (Glycosylation)[40], GPS-SNO (S-nitrosylation)[41], GPS-SUMO (Sumoylation)[42], PrePs (Prenylation)[43], GPS-PAIL (N-acetylation)[44], GPS-Palm (Palmitoylation)[45], GPS-Uber (Ubiquitination)[46], GPS-MSP (Methylation)[47], KprFunc (Propionylation)[48].

### Molecular Dynamics Simulations

Molecular dynamic simulations were used to study the conformational transition of the *VCP* protein when mutations were induced into it. It provides a direct connection between the structure and dynamics via the vast exploration of the conformational energy landscape which is accessible to the protein molecules. The GROMACS software was utilized to investigate *VCP* both in the wild type plus the mutant variants. For this study, GROMOS96 43a1 was used as the forcefield. Considering the dissolvable water surrounding the protein model spc18 was utilized. The framework utilized was the steepest descent approach to minimize the parameters of energy, equilibrated at 310 Kelvin temperature, 1 bar pressure using the canonical ensemble (NVT) (constant number of particles, temperature, and volume) and the (NPT) outfit (constant number of particles, temperature, pressure)[49]. Additionally in MD run parameters simulation time was rose to 50 nsec, MD integrator Leap-frog was used and 5000 frames were used per simulation. Root-mean square deviation (RMSD), Root-mean square fluctuation (RMSF), Radius of gyration (RoG), H-bonds between protein and water, Surface accessible solvent area (SASA), were all calculated and plotted.

## Results

### Protein-Protein Interaction and Disease Pathology of *VCP*

The interactions of the *VCP* protein showed that it majorly interacts with 48 proteins. The subcellular location of the protein was found to be in the Nucleoplasm, cytosol and the predicted location being intracellular. *VCP* was found to have low tissue specificity, it belongs to the protein class of enzymes and transporters. *VCP* has low tissue, and low immune cell specificity too. The pathology of the *VCP* protein was studied to understand its expression in various diseases. The pathology analysis of *VCP* was done for breast cancer, carcinoid, cervical cancer, colorectal cancer, endometrial cancer, glioma, head and neck cancer, liver cancer, lung cancer, lymphoma, melanoma, ovarian cancer, pancreatic cancer, prostate cancer, renal cancer, skin cancer, stomach cancer, testis cancer, thyroid cancer, and urothelial cancer. For all these diseases *VCP* was not prognostic but it was found that renal cancer and endometrial cancer had high expression for *VCP*, and the protein was prognostic for these diseases as well, the p-value for both was found to be less than 0.001. Hence *VCP* can be considered as a prognostic marker for both renal cancer and endometrial cancer[8].

**Figure 1:**
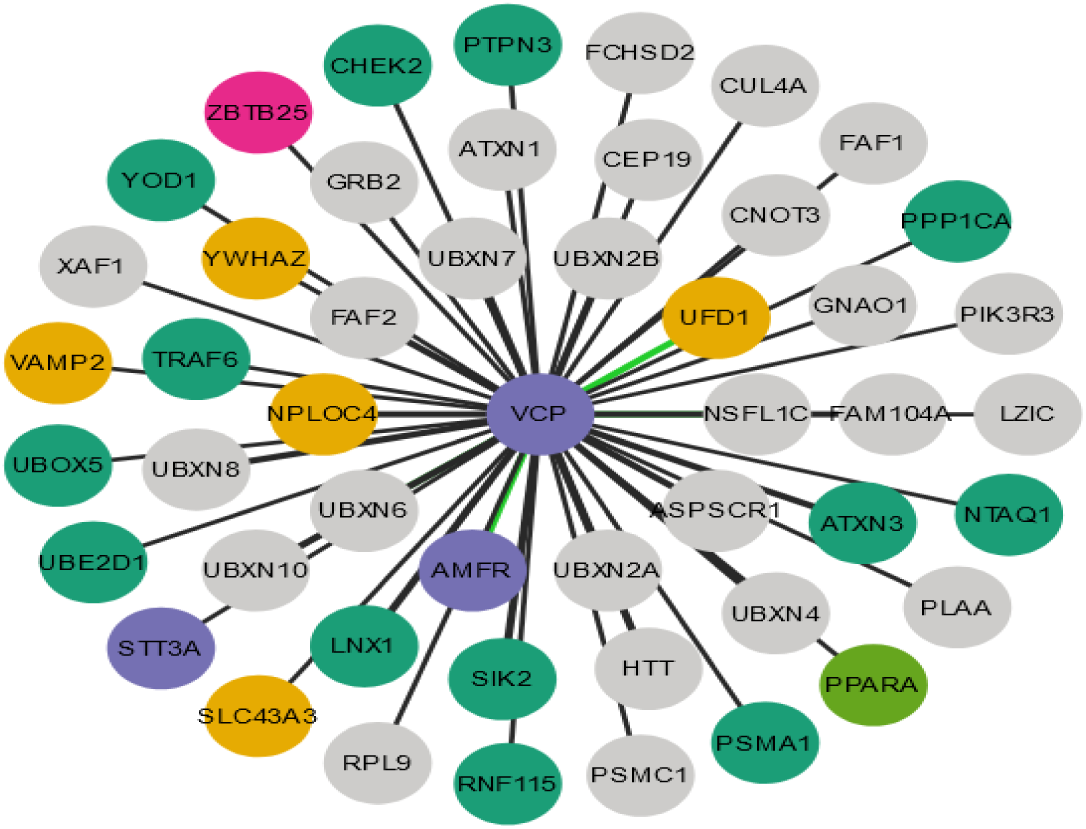
Protein-protein interaction of the *VCP* protein. (colour-yes)

The green edge showed a direct association with *UFD1*. The colour code of the protein depicts the classes of the protein *VCP* interacts with. Dark green being only enzymes, violet for enzymes and transporters, pink being transcription factors, light green for transcription factors and transporters, yellow ochre for only transporters, and grey for others.

**Figure 2:**
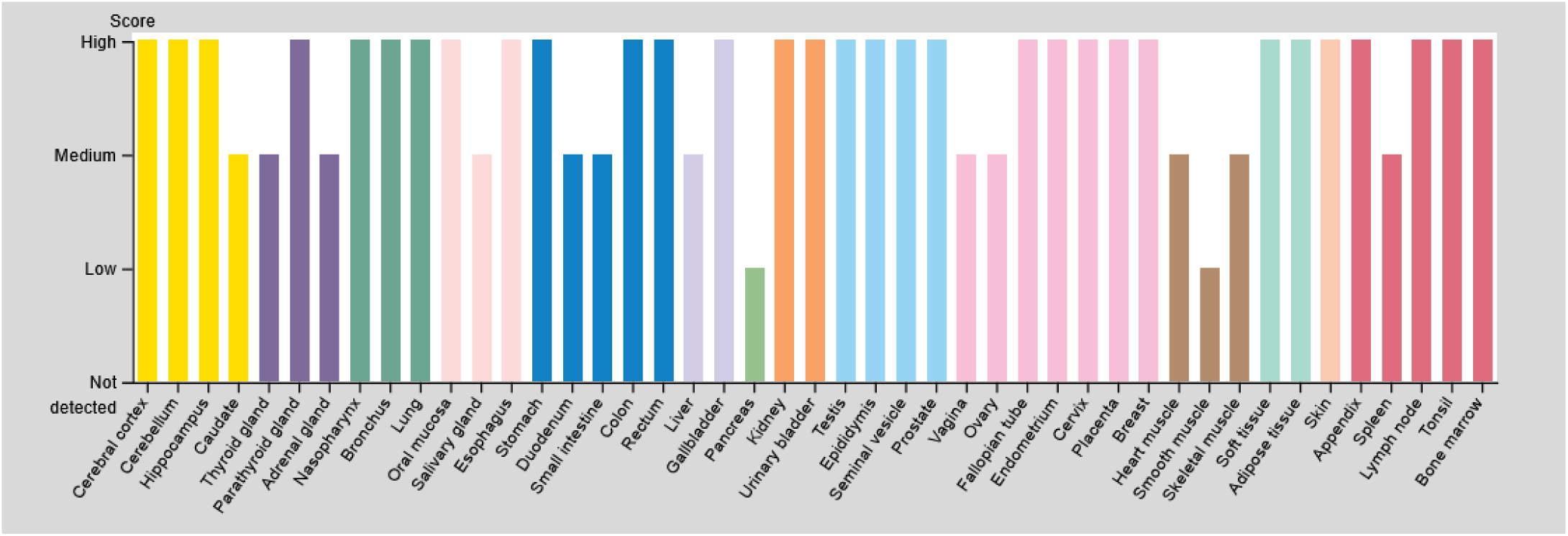
Expression of the *VCP* protein. (colour-yes)

**Figure 3:**
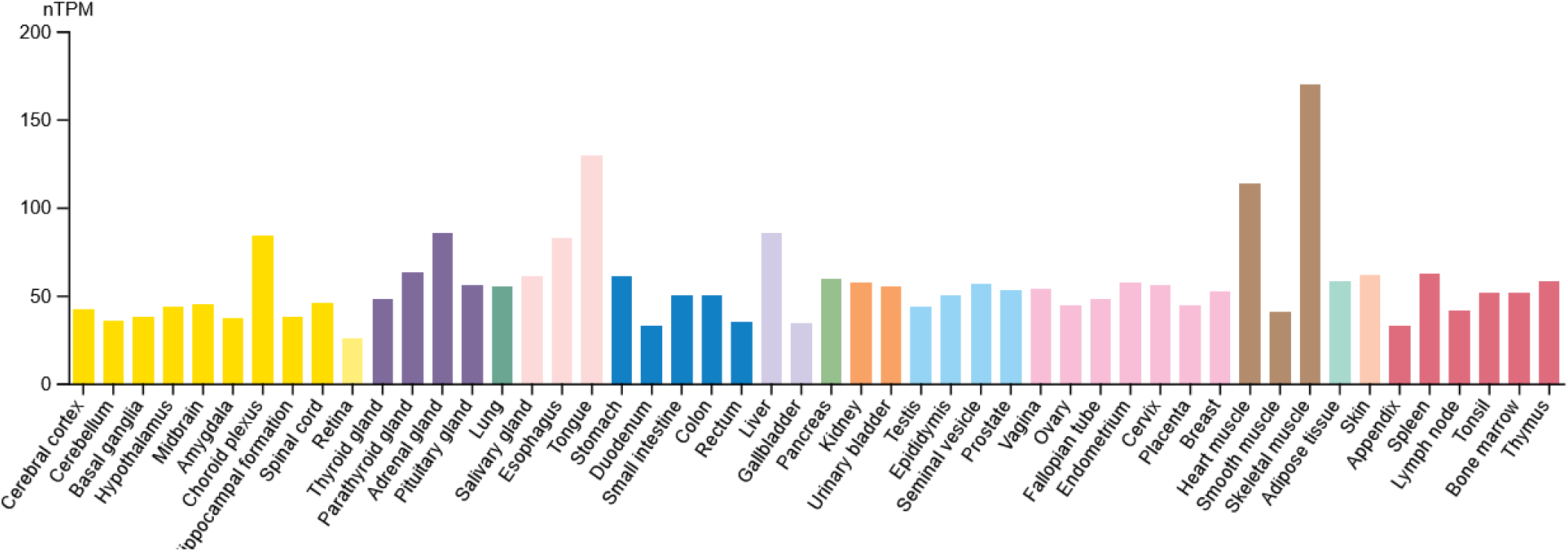
RNA tissue specificity. *VCP* showed to have low tissue specificity. (colour-yes)

### Pathogenicity Prediction

The four variants of *VCP* which fall under the criteria of Pathogenic and likely pathogenic R95C, R95G (rs121909332), and R191P, R191Q (rs121909334) were analysed for pathogenicity using various bioinformatics tools to establish the deleteriousness of the variants and to garner concrete insights to know whether the mutations can lead to fatal diseases. The variant with conflicting interpretation of pathogenicity was also investigated such that A160P (rs1554668805) mutation can fall under a proper, significant class of being deleterious or benign.

**Table 1:** Prediction of pathogenicity of the *VCP* variants.

| rsID | Amino Acid change | SIFT | POLYPHEN2 | PANTHER | SNP&GO | PHDSNP | MetaRNN |
| --- | --- | --- | --- | --- | --- | --- | --- |
|  | (Pathogenic likely Pathogenic) |  |  |  |  |  |  |
| **rs121909332**<br>**283C>T** | R95C | Not tolerated | Probably damaging (Score – 0.993) | Probably damaging (Preservation time – 1629 my) | Disease (Reliability Index -7) | Disease Polymorphism (Reliability Index -5) | 0.7522 |
| **283C>G** | R95G | Tolerated | Probably damaging (Score – 0.999) | Probably damaging (Preservation time – 1629 my) | Disease (Reliability Index -2) | Neutral Polymorphism (Reliability Index -2) | 0.9404 |
| **rs121909334**<br>**572G>C** | R191P | Not tolerated | Probably damaging (Score – 1.0) | Probably damaging (Preservation time – 1629 my) | Disease (Reliability Index -8) | Disease Polymorphism (Reliability Index -4) | 0.9019 |
| **572G>A** | R191Q | Tolerated | Probably damaging (Score – 1.0) | Probably damaging (Preservation time – 1629 my) | Disease (Reliability Index -1) | Neutral Polymorphism (Reliability Index -3) | 0.8911 |
|  | (Conflicting interpretations of pathogenicity) |  |  |  |  |  |  |
| **rs1554668805**<br>**478G>C** | A160P | Tolerated | Benign (Score - 0.002) | Probably damaging (Preservation time – 1039 my) | Disease (Reliability Index -5) | Disease Polymorphism (Reliability Index -0) | 0.5813 |

**Table 2:** The table depicts the Mutpred2 pathogenicity results of the *VCP* variants.

| rsID | Amino Acid change | MutPred2 score | Remarks | Affected PROSITE and ELM Motifs |
| --- | --- | --- | --- | --- |
|  | (Pathogenic likely Pathogenic) |  |  |  |
| **rs121909332** | R95C | 0.429 | - | - |
|  | R95G | 0.448 | - | - |
| **rs121909334** | R191P | 0.912 | - | ELME000002,<br>ELME000100,<br>ELME000108,<br>ELME000155 |
| Molecular mechanisms with P-values $\leq 0.05$ | | | Probability | P-value |
| Gain of Intrinsic disorder |  |  | 0.30 | 0.05 |
| Gain of Loop |  |  | 0.29 | 0.01 |
| Altered Transmembrane protein |  |  | 0.28 | 6.5e-04 |
| Gain of B-factor |  |  | 0.27 | 9.8e-03 |
| Loss of Relative solvent accessibility |  |  | 0.27 | 0.02 |
| Altered Metal binding |  |  | 0.13 | 0.05 |
|  | R191Q | 0.609 | - | ELME000002, ELME000100, ELME000108 |
| Molecular mechanisms with P-values $\leq 0.05$ | | | Probability | P-value |
| Altered Transmembrane protein |  |  | 0.29 | 3.8e-04 |
| Gain of B-factor |  |  | 0.25 | 0.02 |
| Loss of Relative solvent accessibility |  |  | 0.25 | 0.03 |
| **rs1554668805** | A160P | 0.928 | - | ELME000137, PS00008 |
| Molecular mechanisms with P-values $\leq 0.05$ | | | Probability | P-value |
| Altered Ordered interface |  |  | 0.32 | 0.01 |
| Gain of ADP-ribosylation at R159 |  |  | 0.22 | 0.02 |
| Altered Transmembrane protein |  |  | 0.21 | 4.2e-03 |
| Gain of Allosteric site at F163 |  |  | 0.19 | 0.05 |

### Protein Stability Prediction

The stability prediction of the mentioned *VCP* variants was done using various *in silico* tools. The stability prediction of proteins is quintessential to understand the molecular origins and what changes can cause the protein to be unstable, leading to disease phenotype.

**Table 3:** Prediction of protein stability on *VCP* variants.

| rsID | Amino Acid change | Maestro |  | Cupsat (kcal/mol) | mCSM (kcal/mol) | I-mutant (kcal/mol) | SDM | Dynamut (kcal/mol) | MUPro (kcal/mol) | DUET |
| --- | --- | --- | --- | --- | --- | --- | --- | --- | --- | --- |
| Pathogenic likely pathogenic |  | ddG (kcal/mol) | Confidence prediction (kcal/mol) |  |  |  |  |  |  |  |
| **rs121909332** | R95C | 0.199<br>(Destabilizing) | 0.923 | -0.04<br>Destabilizing | -0.92<br>Destabilizing | -1.29<br>Destabilizing | -0.190<br>Destabilizing | -0.006<br>Destabilizing | -1.38<br>Destabilizing | -0.976<br>Destabilizing |
|  | R95G | 0.509<br>(Destabilizing) | 0.870 | 0.35<br>Stabilizing | -1.246<br>Destabilizing | -1.81<br>Destabilizing | 0.100<br>Stabilizing | -0.800<br>Destabilizing | -2.22<br>Destabilizing | -1.161<br>Destabilizing |
| **rs121909334** | R191P | 0.271<br>(Stabilizing) | 0.925 | 0.26<br>Stabilizing | -0.518<br>Destabilizing | -1.40<br>Destabilizing | -1.330<br>Destabilizing | 0.371<br>Stabilizing | -1.22<br>Destabilizing | -0.796<br>Destabilizing |
|  | R191Q | 0.170<br>(Destabilizing) | 0.932 | -0.47<br>Destabilizing | -0.786<br>Destabilizing | -1.31<br>Destabilizing | -0.900<br>Destabilizing | -0.092<br>Destabilizing | -0.52<br>Destabilizing | -0.809<br>Destabilizing |
| (Conflicting interpretations of pathogenicity) |  |  |  |  |  |  |  |  |  |  |
| **rs1554668805** | A160P | -0.110<br>(Stabilizing) | 0.849 | 1.31<br>Stabilizing | -0.696<br>Destabilizing | -2.21<br>Destabilizing | -1.850<br>Destabilizing | -0.112<br>Destabilizing | -1.21<br>Destabilizing | -0.777<br>Destabilizing |

### Evolutionary Conservation

The evolutionary conservation of the protein was checked using the Consurf web server and the data was checked for all the variants. The homologues for the evolutionary conservation were taken from two protein databases UniRef90 and UniProt separately.

**Figure 4:**
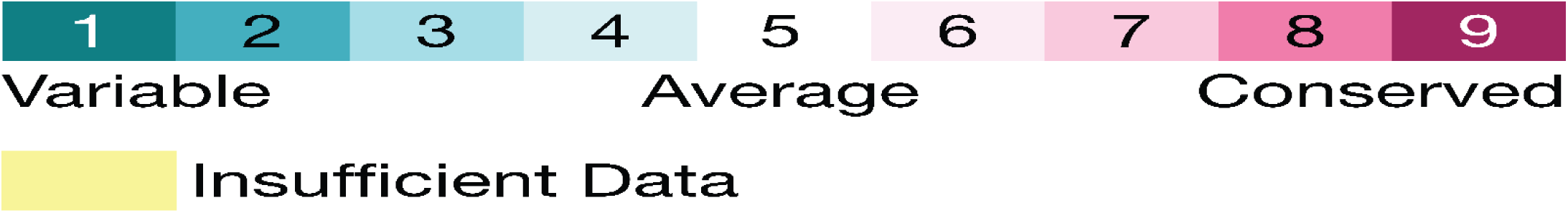
Consurf Evolutionary conservation scale. (colour-yes)

**Table 3:** Evolutionary conservation of *VCP* variants with homologues from UniRef90 database.

| rsID | Amino Acid change | Conservation Score | Localization of residue | Role of residue |
| --- | --- | --- | --- | --- |
|  | Pathogenic likely Pathogenic |  |  |  |
| **rs121909332** | R95C | 5 | Buried | Not available |
|  | R95G | 5 | Buried | Not available |
| **rs121909334** | R191P | 4 | Exposed | Functional |
|  | R191Q | 4 | Exposed | Functional |
|  | (Conflicting interpretations of pathogenicity) |  |  |  |
| **rs1554668805** | A160P | 2 | Buried | Not available |

**Table 4:** Evolutionary conservation of *VCP* variants with homologues from UniProt database.

| rsID | Amino Acid change | Conservation Score | Localization of residue | Role of residue |
| --- | --- | --- | --- | --- |
|  | Pathogenic likely<br>Pathogenic |  |  |  |
| rs121909332 | R95C | 3 | Exposed | Not available |
|  | R95G | 3 | Exposed | Not available |
| rs121909334 | R191P | 3 | Exposed | Not available |
|  | R191Q | 3 | Exposed | Not available |
|  | (Conflicting interpretations of pathogenicity) |  |  |  |
| rs1554668805 | A160P | 1 | Buried | Not available |

### Stereochemical and Physicochemical Analysis

**Figure 5:**
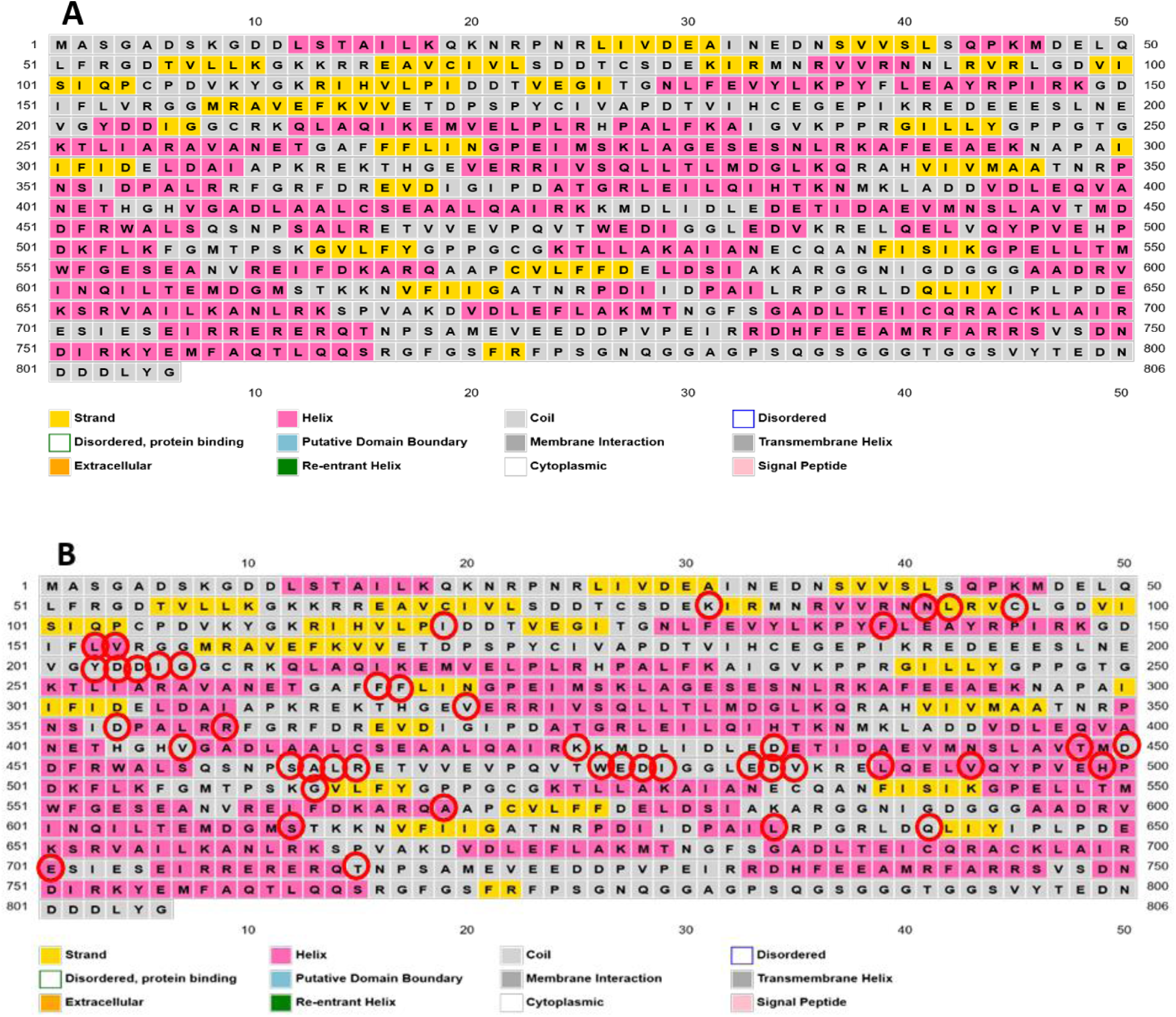
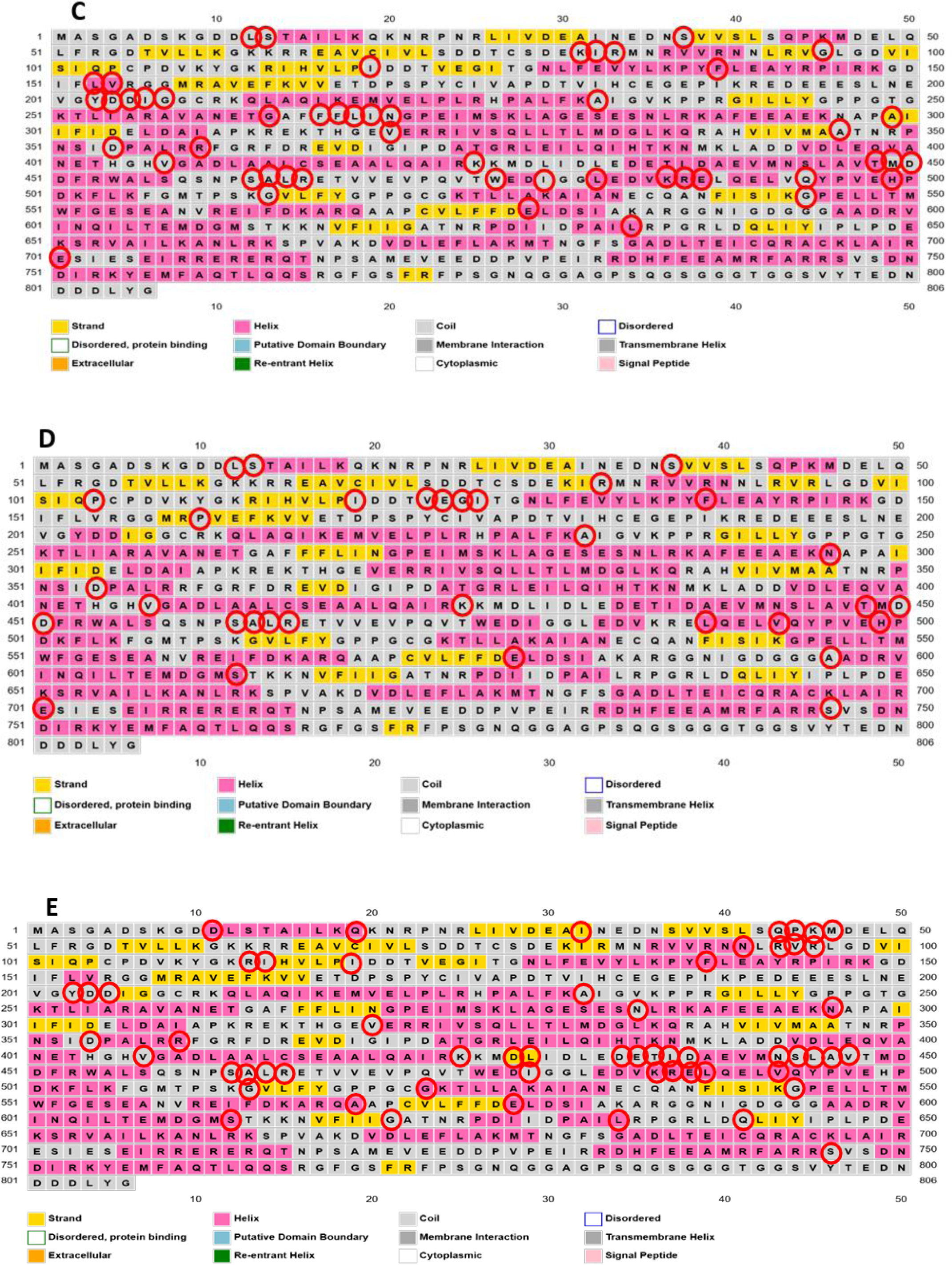
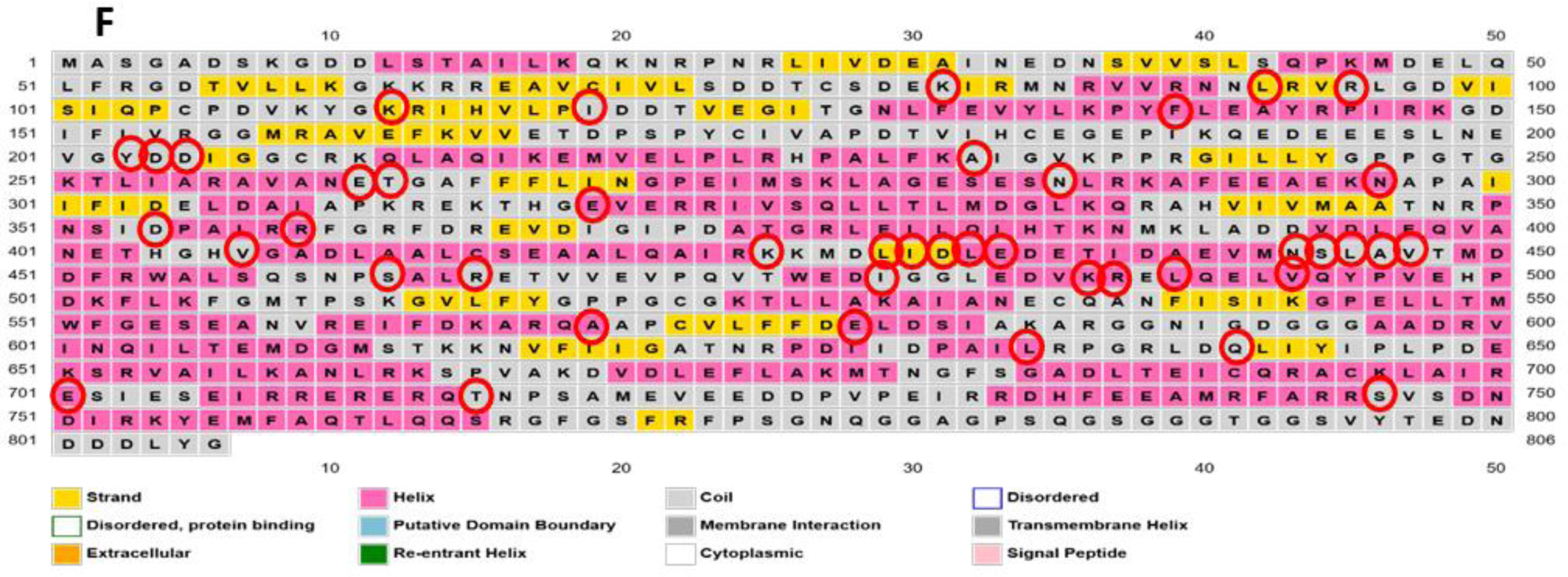
Secondary structure of the *VCP* Wild type and the five variants with the position changes marked A) Wild type B) R95C C) R95G D) A160P E) R191P F) R191Q (colour-yes)

It was observed upon the introduction of the mutations the secondary structure changed significantly. In the wild type, position 95 and 160 were strand with the potential formation of Beta-sheet and, position 191 was Coil. The induction of the mutant variants R95C, R95G, A160P changed the 95^th^ and 160^th^ position to Coil. Upon incorporation of the mutant R191P and R191Q, the 191^st^ position remained unchanged. Apart from the mentioned changes, significant other secondary structure changes were noticed in all the five variants as marked in the figures.

The stereochemical quality of the protein structure was analysed through the Ramachandran plot. Ramachandran plot of wild-type *VCP* protein showed above 95% of the residues in the favoured region. It was noticed that rs121909332 when changed from Arginine to Cysteine (R95C) was under the disallowed region with the coordinates (144.27:0.00). Similarly, on the introduction of G in place of R in the 95^th^ position the residue appeared in the disallowed region with the same coordinates as previous (144.27:0.00). For rs1554668805, when 160^th^ residue changed from Alanine to Proline, the Proline appeared in the verge of allowed and disallowed regions in position (-71.00:-87.80). rs121909334, with Arginine replaced by Proline in the 191^st^ position, led to the residue in the unfavourable region (-71.00:-98.57). Similarly, for R191Q, Glutamine was in the unfavourable region (148.69:125.37).

Certain important scales are used to describe a protein’s function and activity, like Hydropathicity, which has an important role in drug design, biomolecular interactions of the proteins. Hydrophobicity, is an important factor considering the folding of the protein. Polarity is of critical importance considering the structure and function of the protein. Alpha helix, and Beta-Sheet are important parameters for the secondary structure of the protein[12,32]. Hence, to check whether the wild type of the *VCP* protein is getting influenced due to the mutations, all the variants were thoroughly scrutinized to find changes using the protein scales.

**Figure 6:**
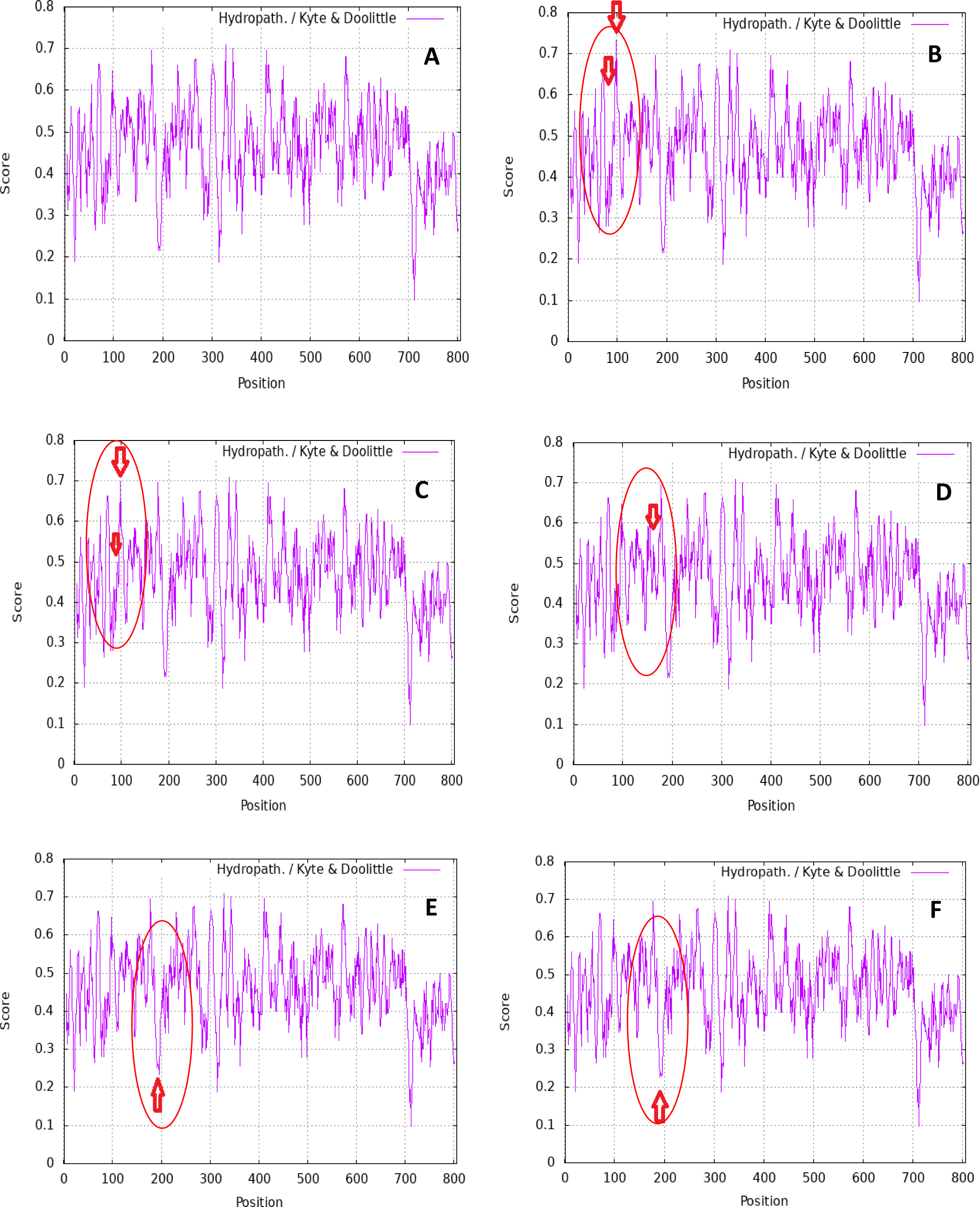
Hydropathicity plots of *VCP* Wild type and the five variants with the position of change marked. A) Wild type B) R95C C) R95G D) A160P E) R191P F) R191Q (colour-no)

**Figure 7:**
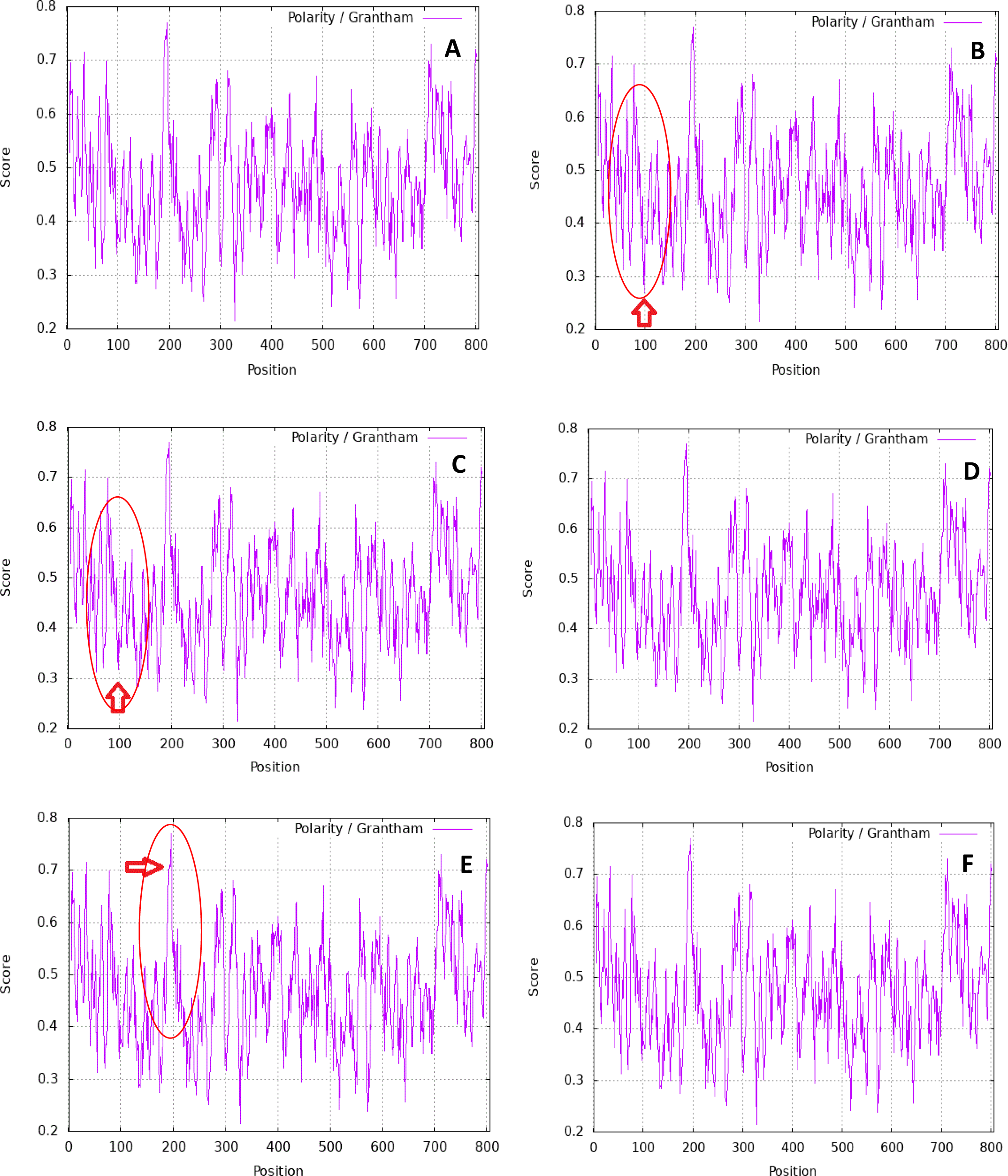
Polarity plots of *VCP* Wild type and the five variants with the position of change marked. A160P & R191Q had no change in Polarity when compared to the Wild type. A) Wild type B) R95C C) R95G D) A160P E) R191P F) R191Q (colour-no)

**Figure 8:**
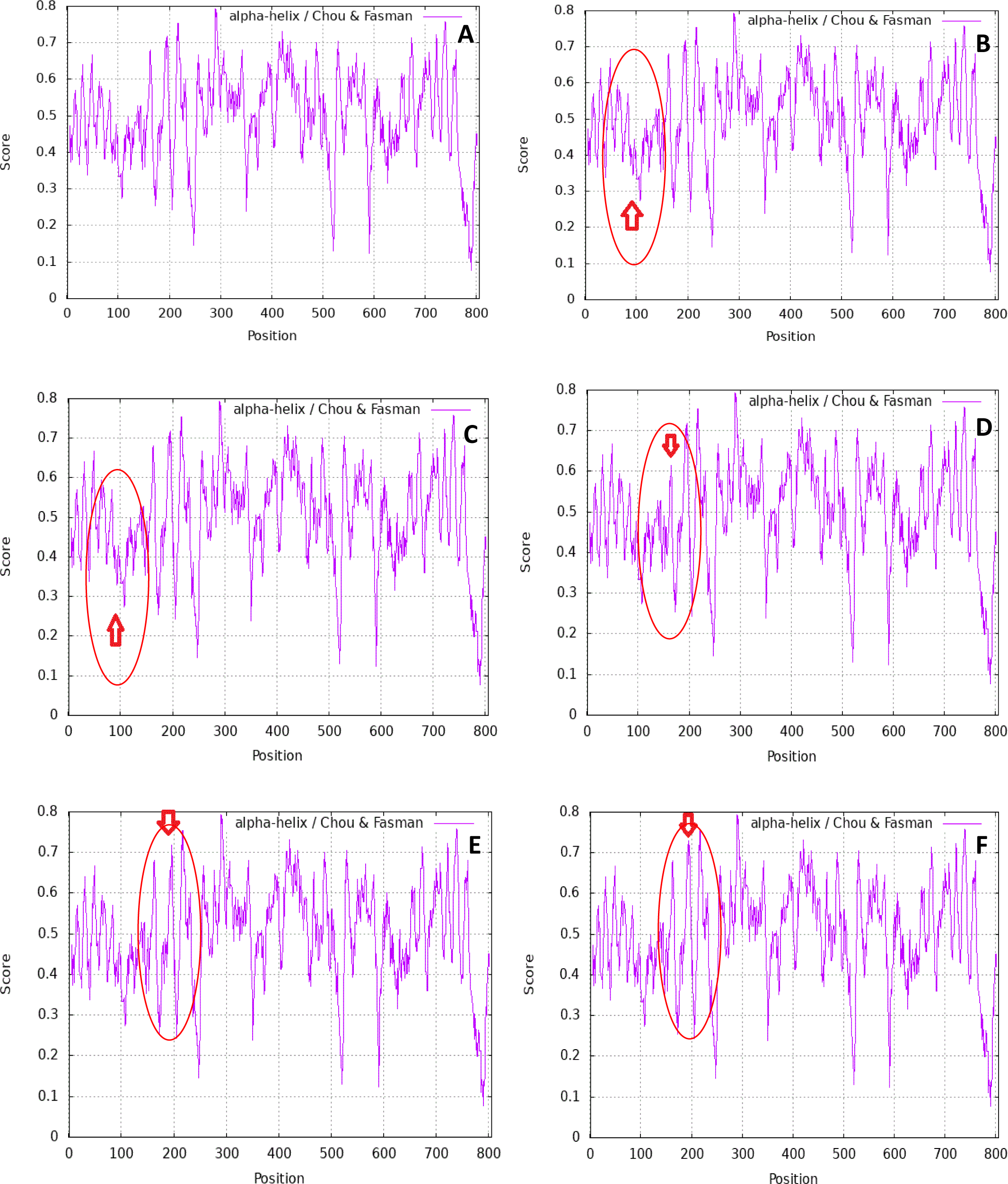
Alpha-Helix plots of *VCP* Wild type and the five variants with the position of change marked A) Wild type B) R95C C) R95G D) A160P E) R191P F) R191Q (colour-no)

**Figure 9:**
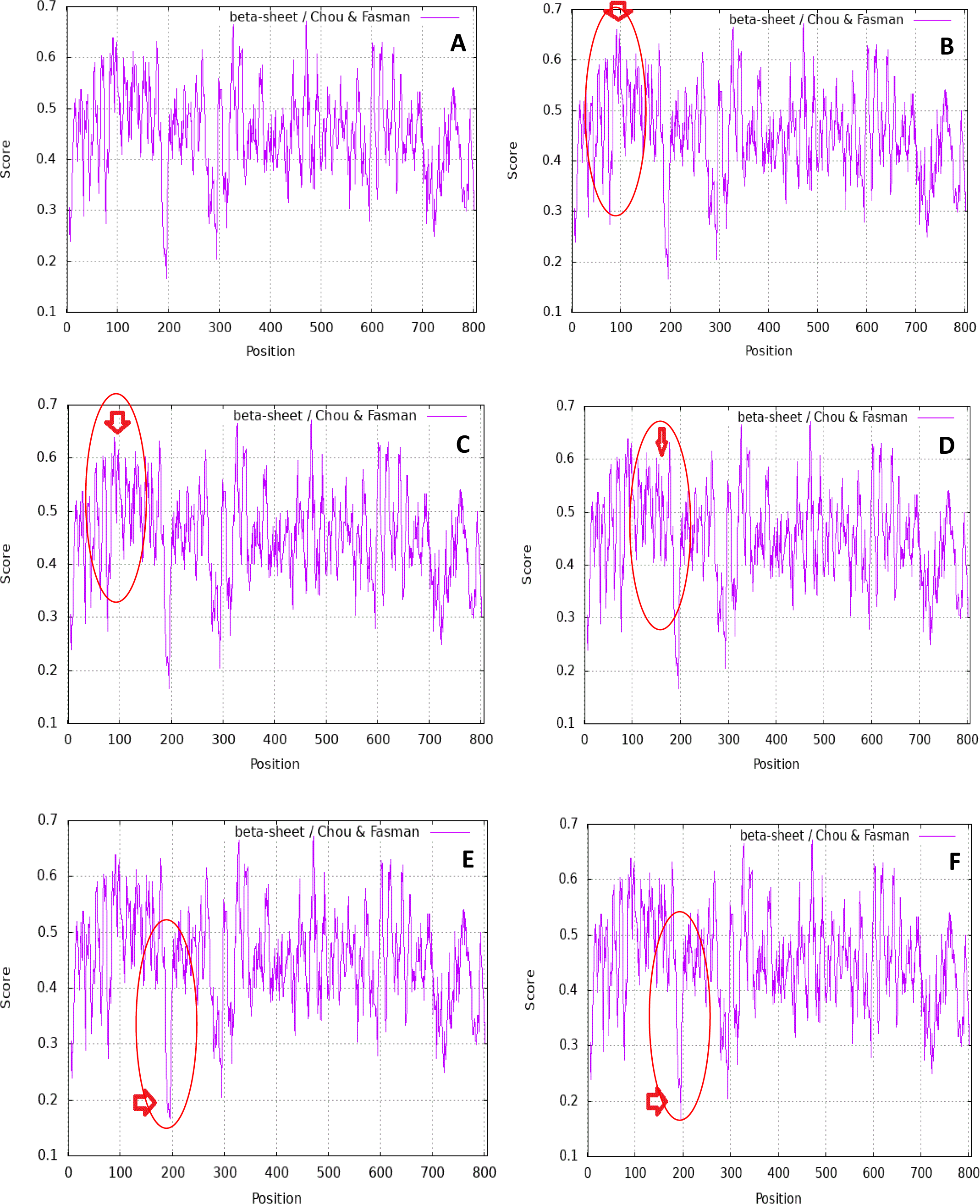
Beta-Sheet plots of *VCP* Wild type and the five variants with the position of change marked. Starting from the top left. A) Wild type B) R95C C) R95G D) A160P E) R191P F) R191Q. (colour-no)

### Genome and OMIM Investigation

The UCSC genome browser was screened for the Valosin-containing protein under Human Assembly GRCH38/hg38 (December 2013 release). *VCP* was observed to be in the position of p13.3 in chromosome 9: 33200001-36300000.

**Figure 10:**
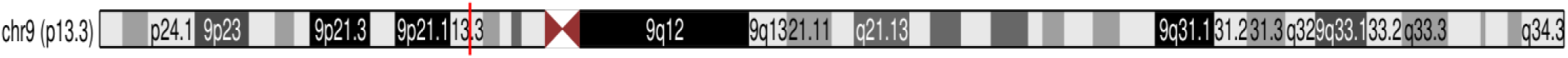
Chromosome 9 (p13.3) (colour-no)

All the rsID of *VCP* were analysed through UCSC and OMIM to understand the extent of severity and the possible associated diseases with it.

**Figure 11:**
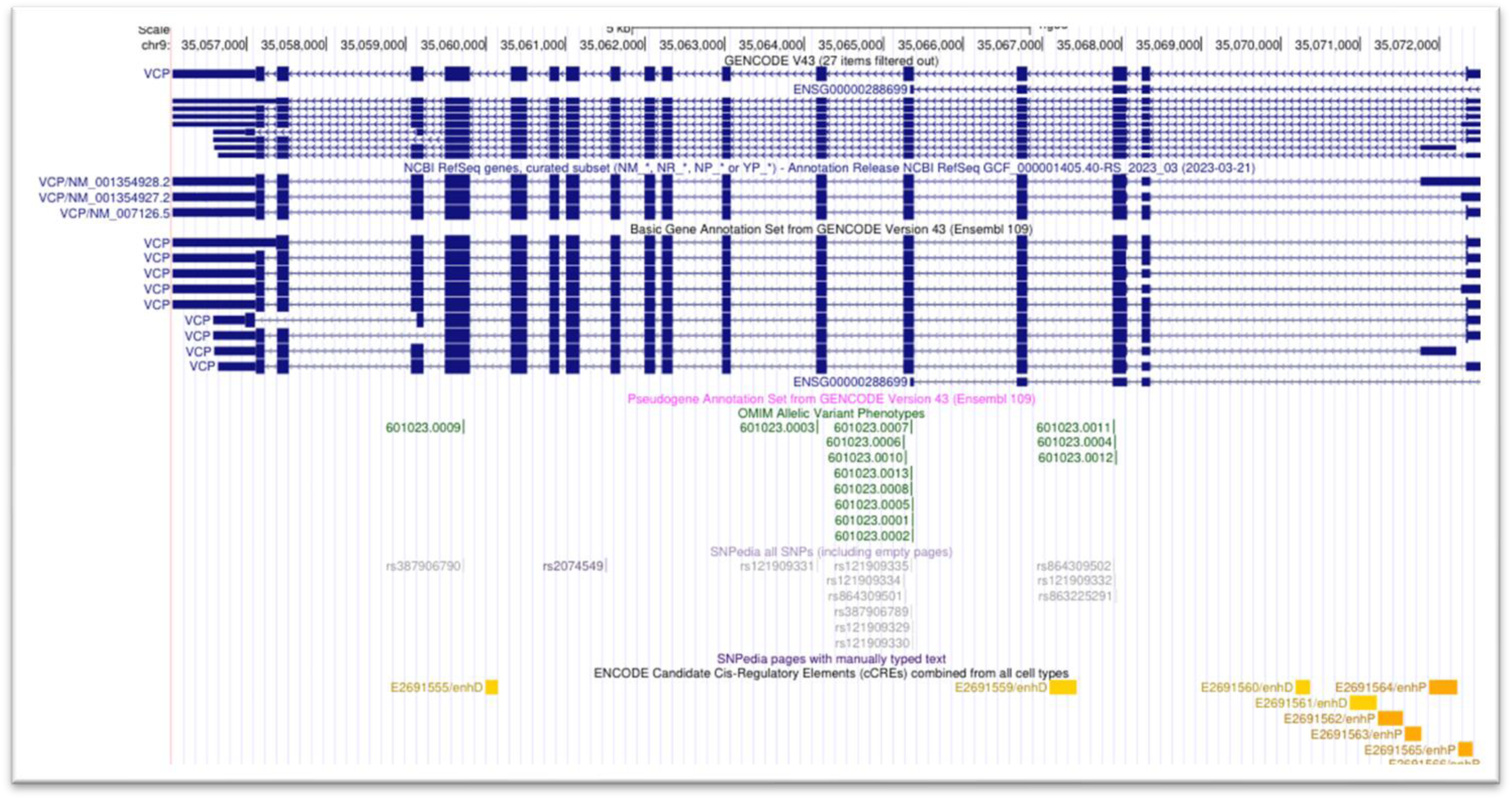
Data from UCSC showing *VCP* under Gencode, NCBI RefSeq, Ensemble along with OMIM variants, SNPedia, and ENCODE Candidate Cis-Regulatory Elements (cCREs). (colour-no)

Upon analysis, rs121909334 (R191Q) with OMIM allelic variant 601023.0006 and position chr9:35065255-35065255 (Band - 9p13.3) was found to be associated with Inclusion Body Myopathy with Early-Onset Paget disease and Frontotemporal Dementia 1. Similarly, rs121909332 (R95G) with OMIM allelic variant 601023.0004 and position chr9:35067910-35067910 (Band - 9p13.3) was also found to be associated with Inclusion Body Myopathy with Early-Onset Paget disease and Frontotemporal Dementia 1.

Under SNPedia and all SNPs, rs121909332 and rs121909334 were found but with no concrete results. There wasn’t enough information about these variants available, this may be due to lack of screening, prove of pathogenicity or disease pathology. Details regarding the variants R95C (rs121909332), R191P (rs121909334), and A160P (rs1554668805) were not found from UCSC combined with the OMIM. In ClinVar variant R95G was deemed as Pathogenic. The Human Assembly GRCH38/hg38 (December 2013 release) showed the ENCODE Candidate Cis-Regulatory Elements (cCREs), which are essential for the regulation of a gene.

### Post Translational Modifications

**Table 5:**
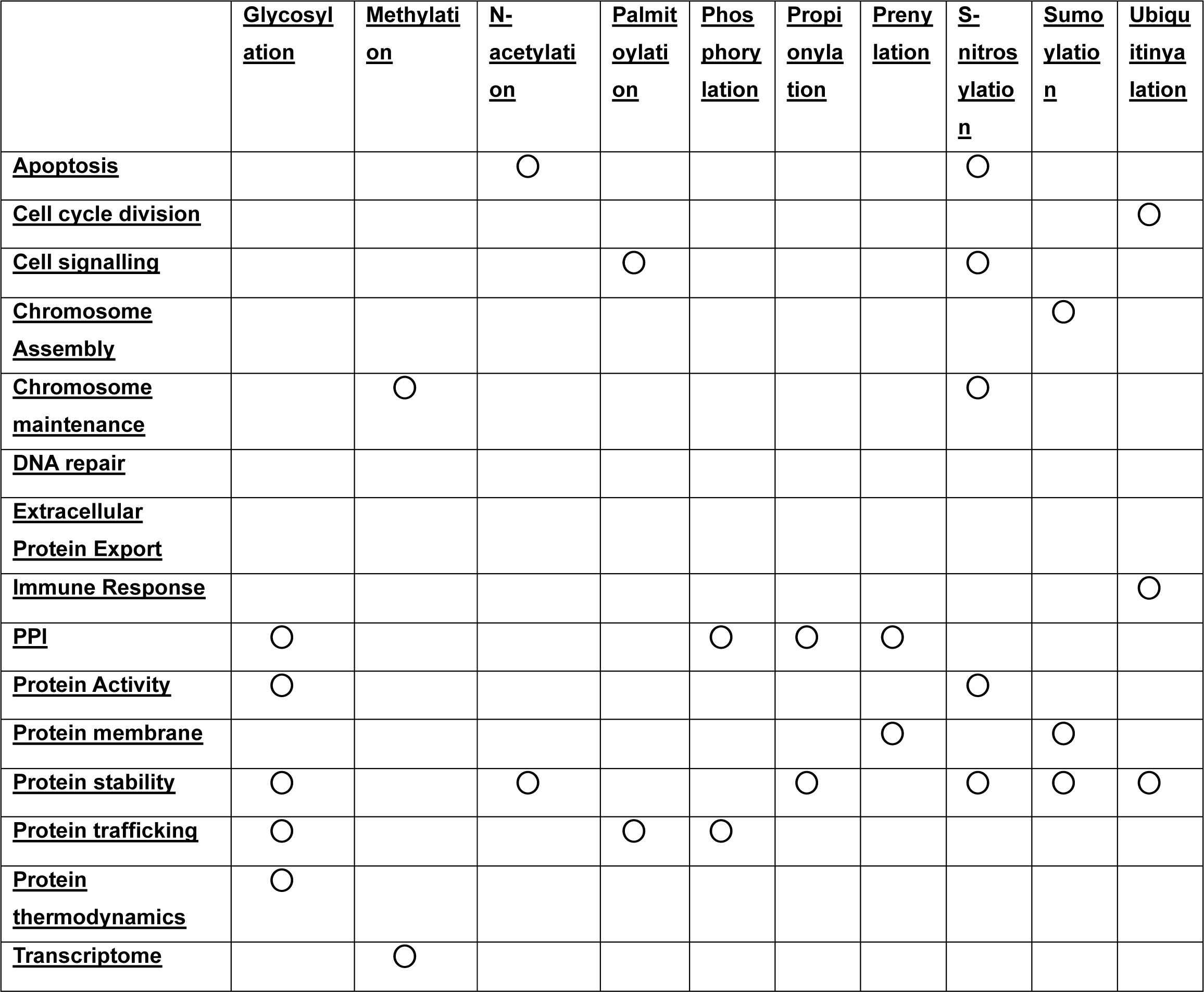
Post-translational modification’s impact on the functional aspects in a cell. The circles depict the functional aspects the PTMs are responsible for.

**Table 6:** Impacts on Post-translational modification due to the *VCP* variants. The variants responsible for changes in the functional aspects are accordingly mentioned in the table.

|  | <u>Glycosylation</u> | <u>Methylation</u> | <u>N-acetylation</u> | <u>Palmitoylation</u> | <u>Phosphorylation</u> | <u>Propionylation</u> | <u>Prenylation</u> | <u>S-nitrosylation</u> | <u>Sumoylation</u> | <u>Ubiquitinylation</u> |
| --- | --- | --- | --- | --- | --- | --- | --- | --- | --- | --- |
| <u>Apoptosis</u> |  |  | ○ |  |  |  |  | R95C |  |  |
| <u>Cell cycle division</u> |  |  |  |  |  |  |  |  |  | ○ |
| <u>Cell signalling</u> |  |  |  | ○ |  |  |  | R95C |  |  |
| <u>Chromosome Assembly</u> |  |  |  |  |  |  |  |  | A160 P, R191 P, R191 Q |  |
| <u>Chromosome maintenance</u> |  | ○ |  |  |  |  |  | R95C |  |  |
| <u>DNA repair</u> |  |  |  |  |  |  |  |  |  |  |
| <u>Extracellular Protein Export</u> |  |  |  |  |  |  |  |  |  |  |
| <u>Immune Response</u> |  |  |  |  |  |  |  |  |  | ○ |
| <u>PPI</u> | ○ |  |  |  | R95C, R95G | ○ | ○ |  |  |  |
| <u>Protein Activity</u> | ○ |  |  |  |  |  |  | R95C |  |  |
| <u>Protein membrane</u> |  |  |  |  |  |  | ○ |  | A160 P, R191 P, R191 Q |  |
| <u>Protein stability</u> | ○ |  | ○ |  |  | ○ |  | R95C | A160 P, R191 P, R191 Q | ○ |
| <u>Protein trafficking</u> | ○ |  |  | ○ | R95C, R95G |  |  |  |  |  |
| <u>Protein thermodynamics</u> | ○ |  |  |  |  |  |  |  |  |  |
| <u>Transcriptome</u> |  | ○ |  |  |  |  |  |  |  |  |

### Molecular Simulations

**Figure 12:**
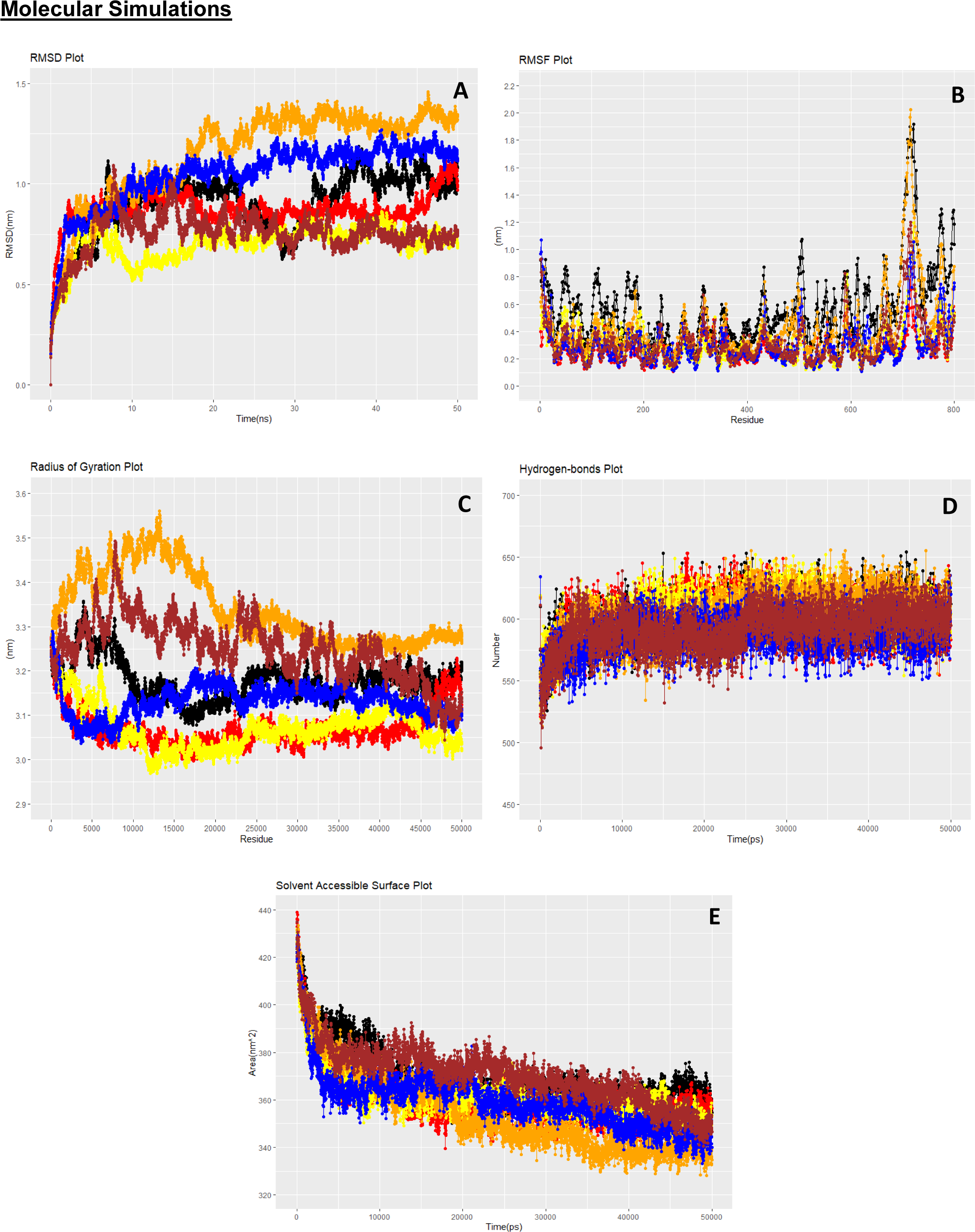
Molecular Dynamic simulations of the *VCP* Wild type and the mutant variants A) RMSD B) RMSF C) Radius of Gyration (RoG) D) Hydrogen bonds E) Solvent Accessible Surface Area (SASA). Colour code: Wild-Black, R95C-Red, R95G-Yellow, A160P-Orange, R191P-Blue, R191Q-Brown. (colour-yes)

## Discussion

Advent of modern sequencing technologies has helped in the investigation of novel variants. Screening and functional analysis of different population cohorts is necessary to differentiate variants as disease causing or benign. This research aimed to functionally investigate genetic variants of the VCP gene which had uncertain significance but have been reported in patients suffering from fatal diseases.

The research was conducted with five variants of *VCP* R95C, R95G, A160P, R191P, R191Q obtained from the NCBI dbsnp database, where R95C, R95G (both rs121909332) and R191P, R191Q (both rs121909334) fell under the criteria of Pathogenic like Pathogenic, whereas A160P (rs1554668805) fell under Conflicting interpretations of Pathogenicity. Upon literature review and analysis, it was found that these variants have been found in individuals with various diseases. rs121909332 was found to be involved in patients suffering from Inclusion Body Myopathy with Early-Onset Paget Disease with or without Frontotemporal Dementia 1, rs121909334 was seen to be involved with Amyotrophic Lateral Sclerosis, Amyotrophic Lateral Sclerosis 14 with or without Frontotemporal Dementia, Charcot-Marie-Tooth disease type 2Y, Inclusion Body Myopathy with Early-Onset Paget Disease with or without Frontotemporal Dementia 1, Dementia, Osteitis Deformans, Inclusion Body Myopathy with Paget’s disease of bone and Frontotemporal Dementia (IBMPFD), Parkinson’s disease[9,10]. But no information could be found for rs1554668805, in regards to its disease relation. The variants were found to be variably conserved, with the lowest conservation score for A160P.

**Figure 13:**
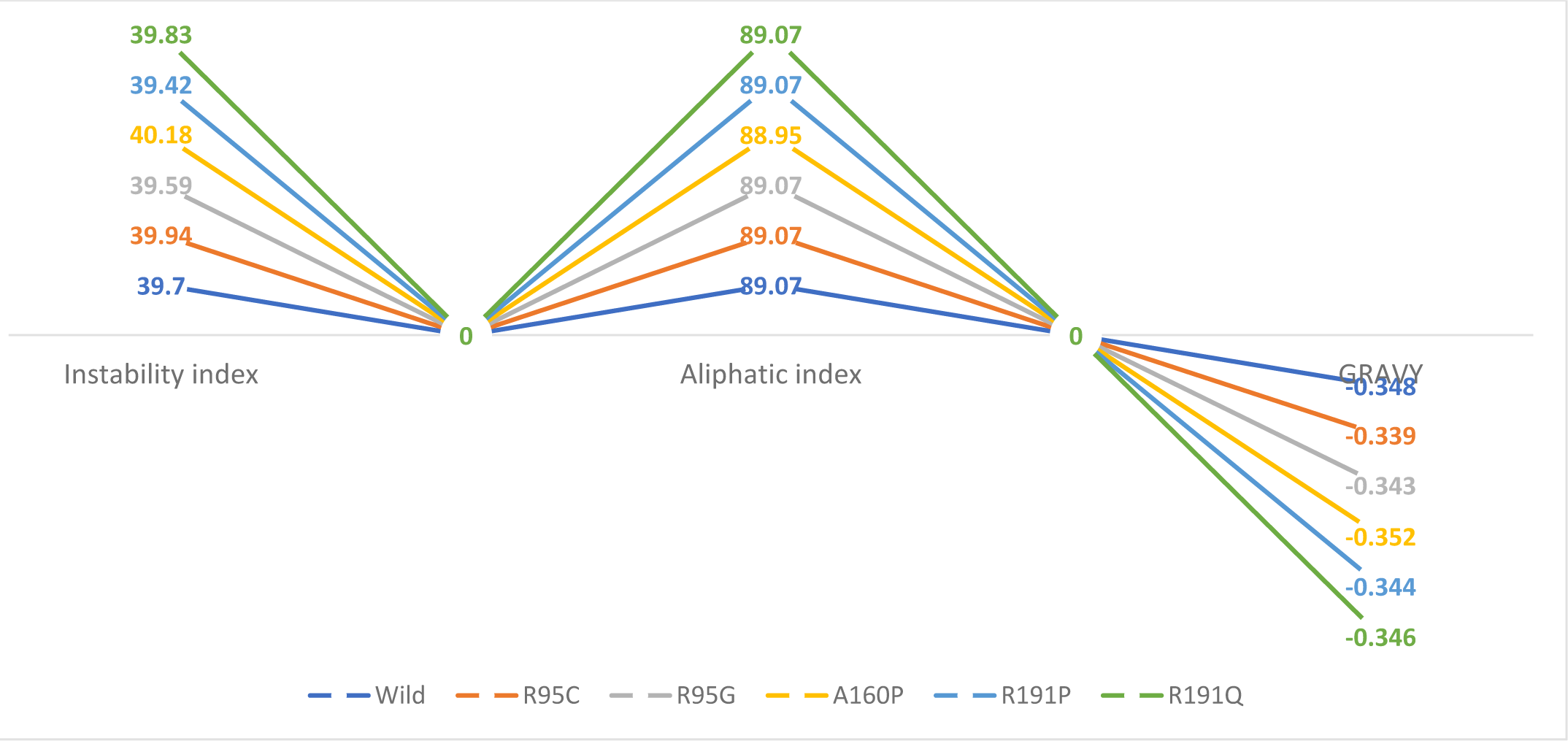
Variation in the Instability Index, Aliphatic Index, and GRAVY values of the mutants in comparison to the Wild type. (colour-yes)

It was seen from the indices above that the values were quite near to the Wild type *VCP* value, in the Instability Index, the highest variation was seen in A160P with a value of 40.18, for the aliphatic index A160P had the lowest value, that is 88.95 within all, in terms of GRAVY as well A160P had the most deviation with the lowest value -0.352.

All amino acids have their geometry, size, charge, hydropathicity, hydrophobicity, Polarity, Alpha-helix, and Beta-Sheet values. The variants often differ in these in comparison to the wild type of the protein. In the case of R95C and R95G, the wild-type residue forms a salt bridge with Aspartic Acid at position 98, and with Glutamic Acid at positions 194, 196, and 200, the difference in charge may affect the interactions made in the general case by the wild type. The backbone of amino acids remains the same always, the changes in properties come due to the change in the side-chain of the mutant from the wild type. As seen for R95C the side chain of the mutant residue is smaller and neutral as compared to the wild-type residue which is larger and positively charged. In the scenario for R191P and R191Q the mutant residue was found to be smaller and neutral than the wild-type residue which was larger and positively charged. The wild type formed a salt bridge with Glutamic Acid in positions 162, 192, 194, and 195. The difference in charge will disturb the interactions made by the wild-type residue. Similarly, for R191Q, the mutant residue was found to be smaller and neutral than the wild-type residue which was larger and positively charged. The variants R95C, R95G, and R191P, were found to be more hydrophobic than the wild type. The wild type formed a salt bridge with Glutamic Acid in positions 162, 192, 194, and 195. The difference in charge will disturb the interactions made by the wild-type residue. For A160P, the mutant residue is bigger than the wild type and was found to be impacting the protein function, unlike the other variants much information couldn’t be found on A160P variation. Considering all the data it can cause possible loss of external interactions of the protein[50]. By Interpro, Gene Ontology, and Broad Gene Ontology, the mutated residue is situated in a domain that is heavily part of the activity of the protein and in contact with other important residues. Due to this a mutation in that position can affect the proper functioning of the protein.

Considering the OMIM analysis, according to Weihl et al. (2006) for Inclusion Body Myopathy with Paget Disease of Bone and Frontotemporal Dementia 1 (IBMPFD1) there was an increase in the conjugates of ubiquitin which were diffused and aggregated and hence the improper function of the endoplasmic reticulum related degradation was observed, additionally, disproportionate Endoplasmic reticulum structure was noticed when cells were transfected with R95G (OMIM ID: 601023.0004)[51]. It was shown by Ju et al. that upon mutations in the VCP gene, the proper clearance of the aggregated proteins was disturbed. R191Q (OMIM ID: 601023.0006) was found in patients suffering from frontotemporal dementia along with/ amyotrophic lateral sclerosis-6 (FTDALS6).

Autosomal dominant inclusion body myopathy with early-onset Paget disease and frontotemporal dementia 1 (IBMPFD1) was identified in 1 out of 13 families due to a transversion of C to G position 283 of the nucleotide VCP gene, eventually led to the substitution of Arginine with Glycine in the 95^th^ position of the protein sequence.

G to C transversion in the 572^nd^ position of the VCP gene was seen in one family out of a total of 13 families screened which led to change in the 191^st^ position to Glutamine from Arginine. R191Q mutation was identified as heterozygous in 4 affected members of the same Italian family who had FTDALS6. The affected individuals in adulthood had phenotypes of ALS, none had symptoms of Paget’s disease and one of the other patients had mild frontotemporal dementia. In addition, phenotypes of IBMPFD were seen in two distinct men aged around fifty, one had Scapuloperoneal weakness while not having any facial involvement, with increased creatine kinase, and the other person had facial weakness, girdle weakness in the shoulder and anterior foreleg, plus had increased creatine kinase by 4 times. Further investigations showed myopathic patterns and dystrophic changes, later mild dysexecutive syndrome was found in one of the patients, but there was no evidence of Paget’s disease. These findings tend to bring a vast spectrum of the disease-associated causes of *VCP* mutations.

MD simulation results of the wild type and mutants are shown (Figure - 12) depicting the RMSD, RMSF, Radius of gyration, hydrogen bonds and Solvent accessible surface area. MD was utilized to understand the protein’s structural and functional behaviour in the environment and provide data on real-time changes that are happening in the variants in comparison to the wild type and thus helped to understand the phenotypic changes which may happen due to differences in stability, conformation, flexibility. When RMSD analysis was used to compare to Wild type, R95C, R95G, and R191Q were stable, high fluctuation was seen in A160P, and moderate fluctuation in R191P. From the RMSF graph, all the variants looked to have lower flexibility than the wild type only A160P was comparably seen to have higher flexibility than the rest of the variants. The gain or loss of flexibility may directly impact the protein’s functioning. RoG determines the compaction of the proteins, R95C, and R95G were more compact than wild unlike A160P which showed the lowest compaction out of all, and R191Q even had lower compaction than wild type. R191P was seen to be almost similar to the wild. The high value of RoG suggests high flexibility and even less stability. Hydrogen bonds contribute favourably to protein’s stability, R191P had lower average H-bonds than wild, R95G, A160P, R95C, and R191Q. SASA predicts the solvent environment accessibility of the protein, only R191Q showed a little higher solvent accessibility than the wild compared to the other variants which showed lower solvent accessibility than the wild with A160P being the lowest of all. SASA is of intense interest as it has direct roles in functional studies and protein folding.

**Figure 14:**
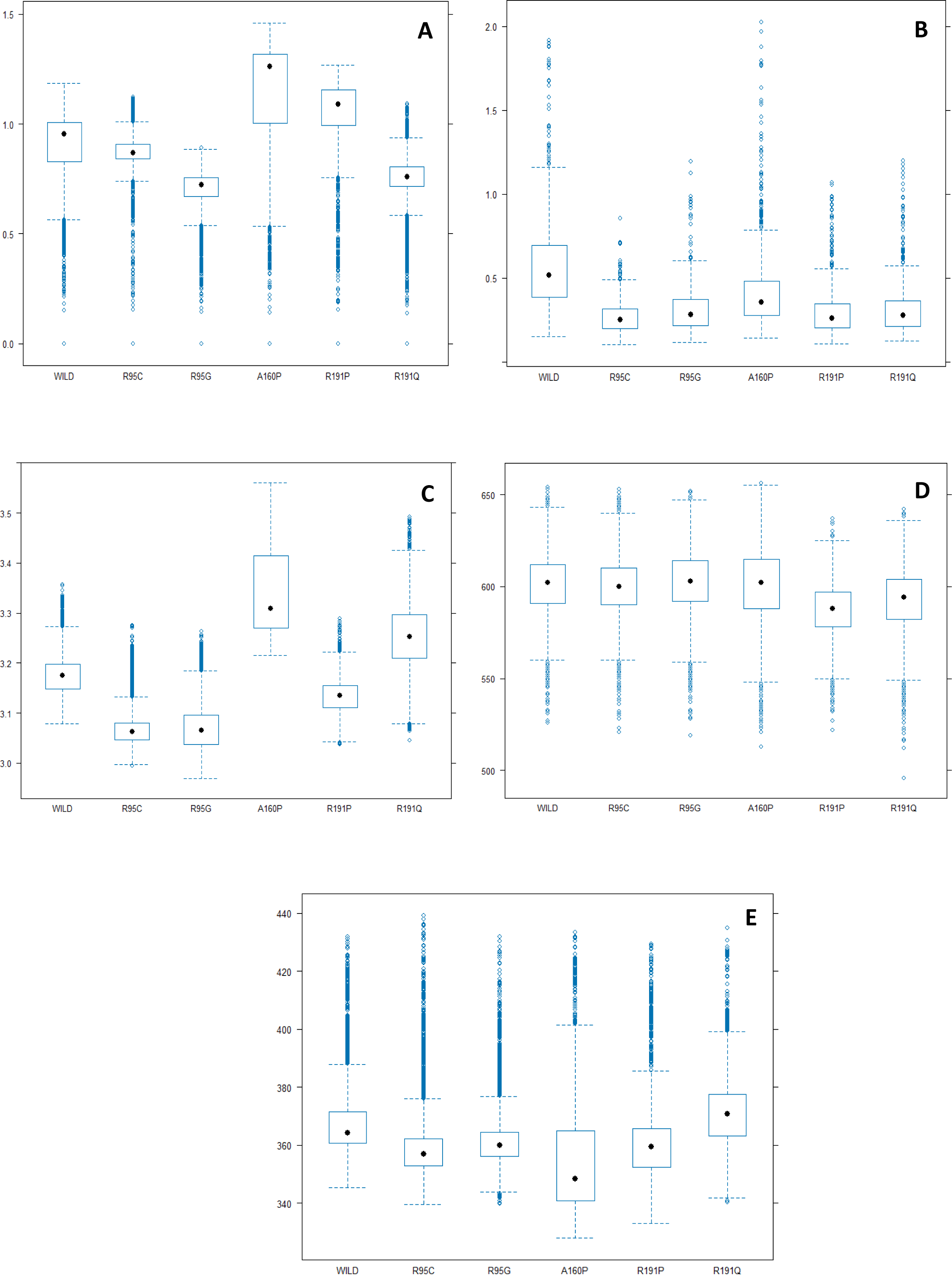
Detailed visualization of the Molecular dynamics data of *VCP* wild type and the variants. A) RMSD B) RMSF C) Radius of Gyration D) Hydrogen bonds E) Solvent Accessible Surface Area (SASA). (colour-no)

The analysis of post-translational modifications showed three parameters being affected due to R95C, R95G (affects phosphorylation), R95C (affects S-nitrosylation), A160P, R191P, R191Q (affects Sumoylation), respectively. It should be noted that all the post-translational modifications were checked against the *VCP* wild type. The changes in Phosphorylation seen in the variants R95C, R95G were found to be below the threshold, hence the changes mayn’t affect the PTM of *VCP* to a very large extent. The regions impacted by S-nitrosylation in the *VCP* wild type were found to be in the amino acid positions 77, 209, and 695. Similarly, for Sumoylation the sites impacted were 8, 136, 164, 486, and 754 in the wild-type *VCP* variant. Consecutively, the same was checked for N-acetylation which showed changes in 8, 236, and 668, in Methylation change was in the amino acid number 8. Propionylation showed its impact on amino acid sites 60, 109, 502, and 677 in the *VCP* variant respectively. Hence, all the variants of *VCP* were found to be impacting any one or the other PTM out of a total of ten being considered in this study. The structure and the function therefore gets impacted due to the variants of *VCP*, affecting the biological processes. The variants may eminently regulate the functioning due to their impact on the Post-translational modifications.

**Figure 15:**
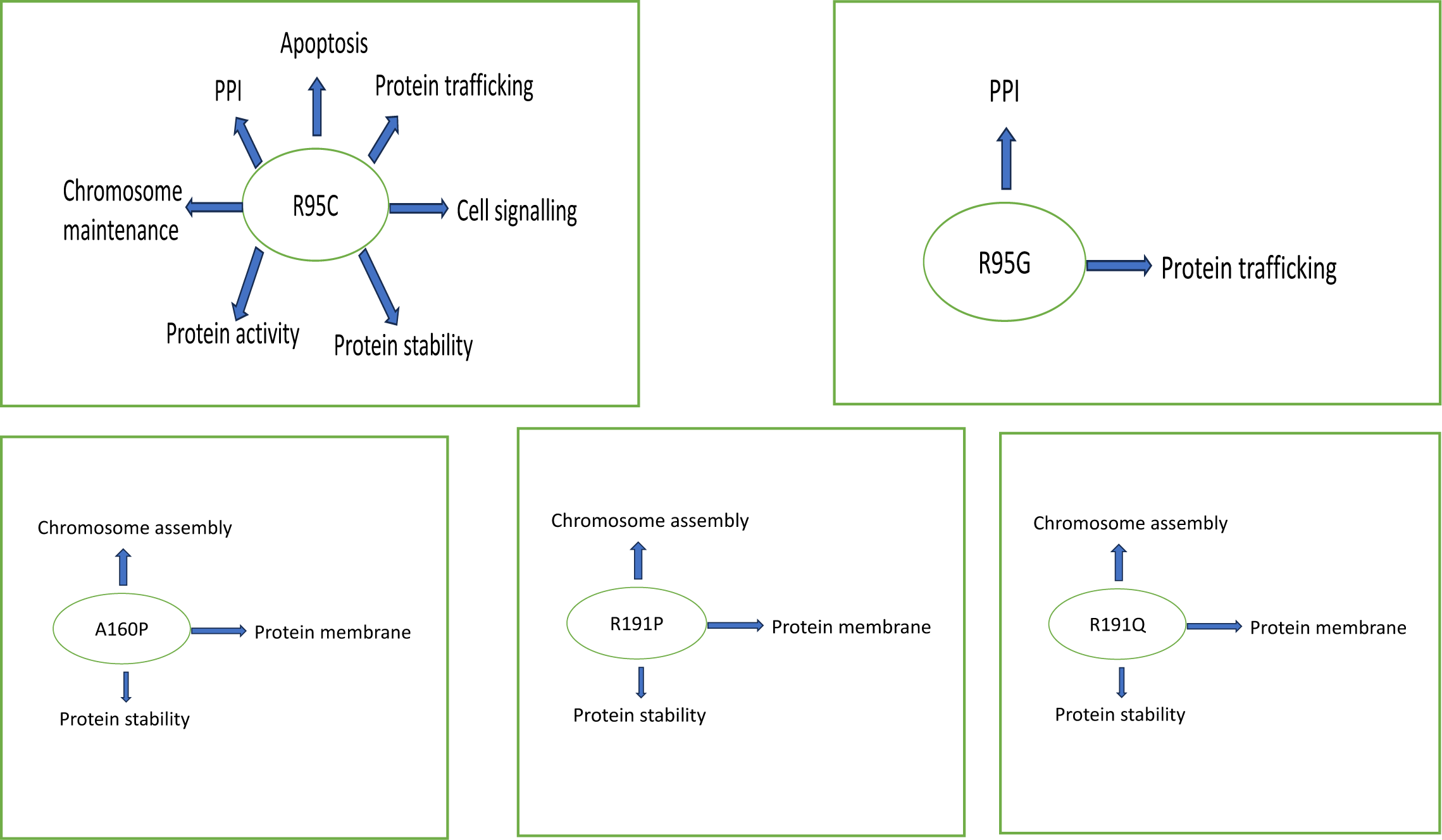
All the *VCP* variants responsible for changes in the various functional domains of the Post-Translational modifications. (colour-no)

**Figure 16:**
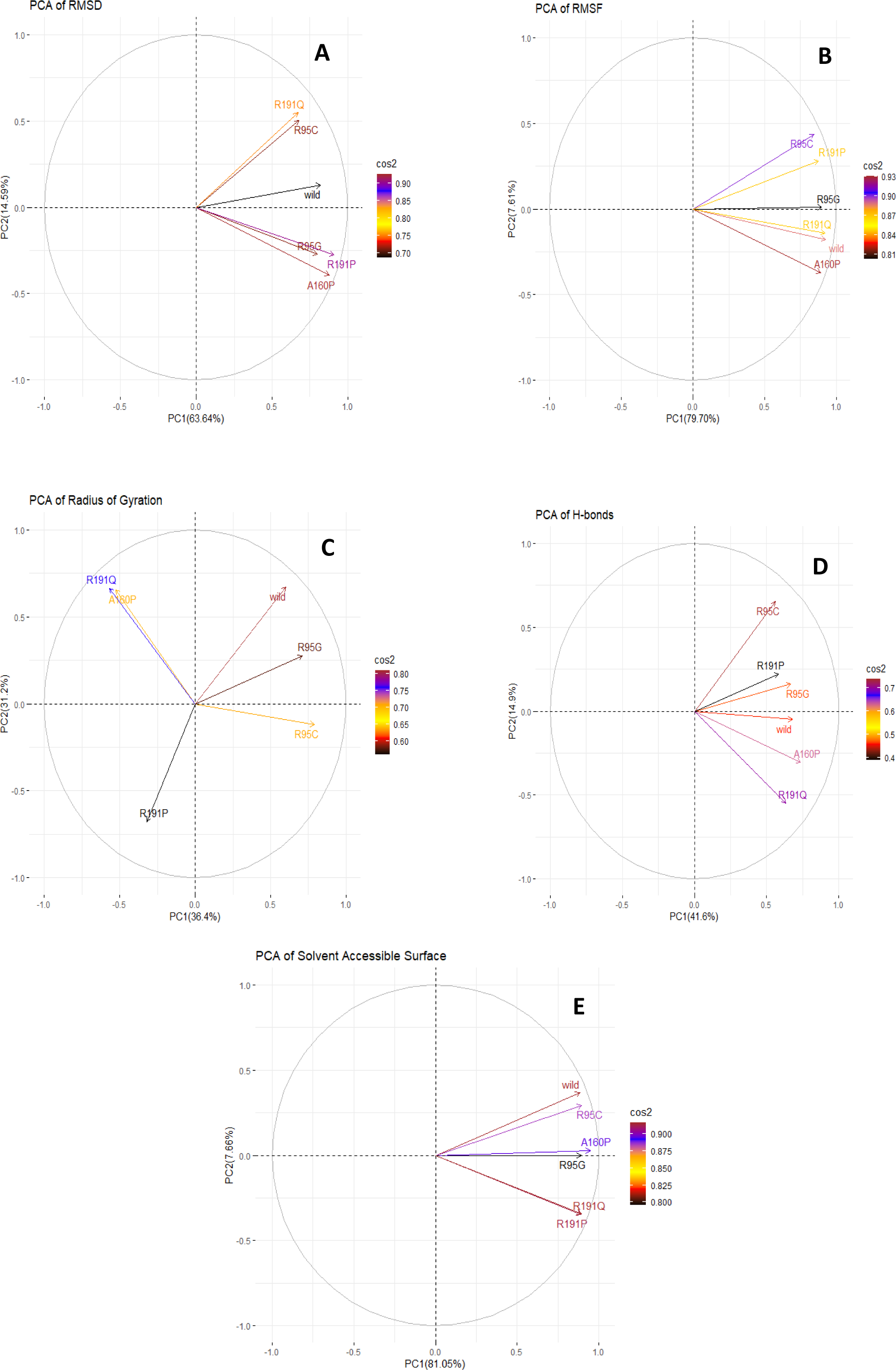
Principal Component Analysis (PCA) of all the *VCP* variants. A) RMSD B) RMSF C) Radius of Gyration D) Hydrogen Bonds E) Solvent Accessible Surface Area (SASA). (colour-yes)

PCA analysis was used to visualize the correlation of the variants against the wild type. PCA makes it easy to visualize complex data. RoG showed the highest negative correlation with R191P making an angle near to 180 degrees against the wild. A detailed explanation of PCA can be found in[52].

## Conclusion

The *in silico* functional investigation of the five missense mutations of *VCP* revealed them to be mildly disease causing. Variants R95G (rs121909332), and R191Q (rs121909334) have heavy evidence against multiple reported diseases. More genetic screening of patients is required for the other three variants. All the five variants showed to destabilize the wild type protein structure. The stereochemical and physicochemical parameters showed significant fluctuations in the mutant proteins. Post-translational modifications showed nine cellular parameters being affected out of a total of fifteen with highest being R95C (rs121909332), which is affecting seven cellular parameters. MD simulations of RMSD, RMSF, RoG, Hydrogen bonds and Solvent accessibility further validated the malfunctioning of the variants. PCA of the MD simulation parameters illustrated the positive and negative correlation of the variants upon comparison with the wild type. This analysis can be referred in wet lab functional study to characterize the significance of these variants and the potential use of these variants as biomarkers in fatal diseases.

## Abbreviations

GRAVY: Grand Average of Hydropathy
RMSD: Root Mean Square Deviation
RMSF: Root Mean Square Fluctuation
RoG: Radius of Gyration
H-Bonds: Hydrogen bonds
SASA: Solvent Accessible Surface Area
IBMPFD: Inclusion Body Myopathy with Paget’s disease of bone and Frontotemporal Dementia
PCA: Principal Component Analysis

## CRediT authorship contribution statement

All work regarding this study was done by Tathagata Das, Saileyee Roy Chowdhury and Professor Parimal Das. The draft of the manuscript was written by Tathagata Das and was checked by Professor Parimal Das. All authors commented on previous versions of the manuscript. All authors read and approved the final manuscript.

## Ethical Approval

Not applicable

## Acknowledgement

Banaras Hindu University, Varanasi, India for providing with internet services. IIT (ISM) Dhanbad, India for allowing to carry out research offsite and AICTE, for providing scholarship to the first author.

